# Local light signalling at the leaf tip drives remote differential petiole growth through auxin-gibberellin dynamics

**DOI:** 10.1101/2022.02.25.481815

**Authors:** Jesse J. Küpers, Basten L. Snoek, Lisa Oskam, Chrysoula K. Pantazopoulou, Sanne E. A. Matton, Emilie Reinen, Che-Yang Liao, Eline D.C. Eggermont, Harold Weekamp, Wouter Kohlen, Dolf Weijers, Ronald Pierik

**Affiliations:** Plant Ecophysiology, dept. Biology, Utrecht University, Padualaan 8, 3584 CH Utrecht, the Netherlands; Theoretical Biology and Bioinformatics, dept. Biology, Utrecht University, Padualaan 8, 3584 CH Utrecht, the Netherlands; Laboratory of Biochemistry, Wageningen University, Stippeneng 4, 6708 WE Wageningen, The Netherlands; Laboratory for Molecular Biology, Wageningen University, Droevendaalsesteeg 1, 6708 PB Wageningen, Netherlands

**Author notes:** Gadeta B.V., Yalelaan 62, 3584 CM Utrecht, the Netherlands. These authors contributed equally to this manuscript (Basten L. Snoek, Lisa Oskam, Chrysoula K. Pantazopoulou).

## Abstract

Although plants are immobile, many of their organs are flexible to move in response to environmental cues. In dense vegetation plants detect neighbours through far-red light perception with their leaf tip. They respond remotely, with asymmetrical growth between the abaxial and adaxial sides of the leafstalk, the petiole. This results in upward movement that brings the leaf blades into better lit zones of the canopy. The plant hormone auxin is required for this response, but it is not understood how non-differential leaf tip-derived auxin can remotely regulate movement. Here we show that remote light signalling promotes auxin accumulation in the abaxial petiole by reinforcing an intrinsic auxin transport directionality. In the petiole, auxin elicits a response of both auxin as well as a second growth promoter; gibberellin. We show that this dual regulation is necessary for hyponastic leaf movement in response to light. Our results reveal how plants can spatially relay information about neighbour proximity from their sensory leaf tips to the petiole base, thus driving adaptive growth.

## Introduction

In dense vegetation, plants adapt their growth to actively compete for light with their neighbours. However, light distribution in vegetation is very heterogeneous and different plant parts will therefore receive different light intensities and density cues ^1^. Plants use intricate mechanisms of signal transfer between plant parts in order to respond adequately to this heterogeneous information ^2–4^, but these mechanisms are still poorly understood. In *Arabidopsis*, adaptive shade avoidance responses to neighbours include hypocotyl elongation in seedlings and petiole elongation and upward leaf movement (hyponasty) in adult plants ^5^. Although adaptive for the individual plant, shade avoidance responses reduce productivity of dense monocultures ^1,6,7^. To accurately evaluate the competitive threat in their environment, plants use phytochrome (phy) photoreceptors to monitor the ratio of red (R) to far-red (FR) light (R/FR) ^8^. In shade, the R/FR is low due to specific absorption of R light by leaves to power photosynthesis. But even before actual shading occurs, reflected FR-enriched light from neighbouring leaves will reduce the R/FR and provide an early neighbour proximity signal that precedes light competition ^9^. FR-enriched light will induce a tissue-specific growth response in *Arabidopsis* leaves depending on the site of perception ^10,11^. FR-enrichment at the petiole locally stimulates petiole elongation while FR-enrichment at the leaf tip (FRtip) results in petiole hyponasty. The spatial separation between FR-induced petiole elongation and hyponasty allows the plant to adjust its growth to optimally respond to either self-shading or neighbour competition ^11^. In FRtip-induced petiole hyponasty there is spatial separation between the leaf tip as the sensory organ and the petiole base as responding organ ^10,11^. Moreover, petiole hyponasty typically requires differential growth rates between the abaxial (bottom) and adaxial (top) sides of the petiole ^12^. This growth response thus provides a study system to unravel how remote light signalling regulates distal and differential growth without local light signalling in the tissue displaying the growth response. We previously established that petiole hyponasty in response to FRtip occurs via local inactivation of phyB in the leaf tip, which typically results in activation of the PHYTOCHROME INTERACTING FACTOR (PIF) bHLH transcription factors ^10,11^. Active PIFs then enhance expression of *YUCCA* (*YUC*) genes that encode the YUC enzymes required for auxin biosynthesis ^13–15^. The auxin that is produced in the leaf tip subsequently stimulates petiole hyponasty. This regulatory network also seems to drive seedling hypocotyl elongation responses upon detection of FR enrichment in the cotyledons ^16,17^.

So far, it remained unclear how the auxin signal that comes from the remote leaf tip directs differential growth and petiole hyponasty. Using tissue-specific time-series RNA-sequencing, we show that neighbour detection in the leaf tip results in unique transcript profiles in the leaf tip as well as the abaxial and adaxial petiole. Leaf tip-derived auxin is specifically transported towards the abaxial petiole to locally enhance gene expression and ultimately cell elongation. Besides auxin, we identify roles for gibberellin (GA) and PIFs in the responding petiole and suggest side-specific activation of the growth-promoting BRASSINAZOLE RESISTANT 1 (BZR1) - AUXIN RESPONSE FACTOR 6 (ARF6) - PIF4 / DELLA (BAP/D) transcription factor module. This study reveals how plants use targeted long-distance auxin signalling to adapt their growth to competitive environments.

## Results

### Characterizing the kinetics and localisation of leaf tip FR light-induced hyponasty and gene expression

Neighbour detection through FR light in the distal leaf tip (FRtip) leads to petiole hyponasty which first becomes visible ~4 hours after start of treatment (Figures 1A and S1A, Video S1). We measured epidermal cell length in the petiole and found that FRtip specifically enhances epidermal cell elongation in the proximal two-thirds of the abaxial petiole (Figure 1B). Considering the previously identified important role of auxin in FRtip induced petiole hyponasty we studied the expression of auxin response genes in the petiole upon FRtip. Indeed, the auxin-responsive transcripts of *IAA29* and *ACS4* were induced in the proximal petiole within 100 minutes of FRtip while the shade marker transcript *PIL1* was unaffected in the non-FR-exposed petiole (Figure S1B). In order to get more insight in the spatial regulation of differential gene expression and petiole growth by FR signalling in the leaf tip, we decided to separately harvest the leaf tip and the separated abaxial and adaxial sides of the proximal two-thirds of the petiole in white light (WL) and FRtip (Figure 1C). To capture the early transcriptional response, we harvested at twenty-minute intervals ranging from 60 minutes (40 minutes for the leaf tip) to 180 minutes of treatment as well as at a 300 minute timepoint (Figure 1D).

**Figure 1.**
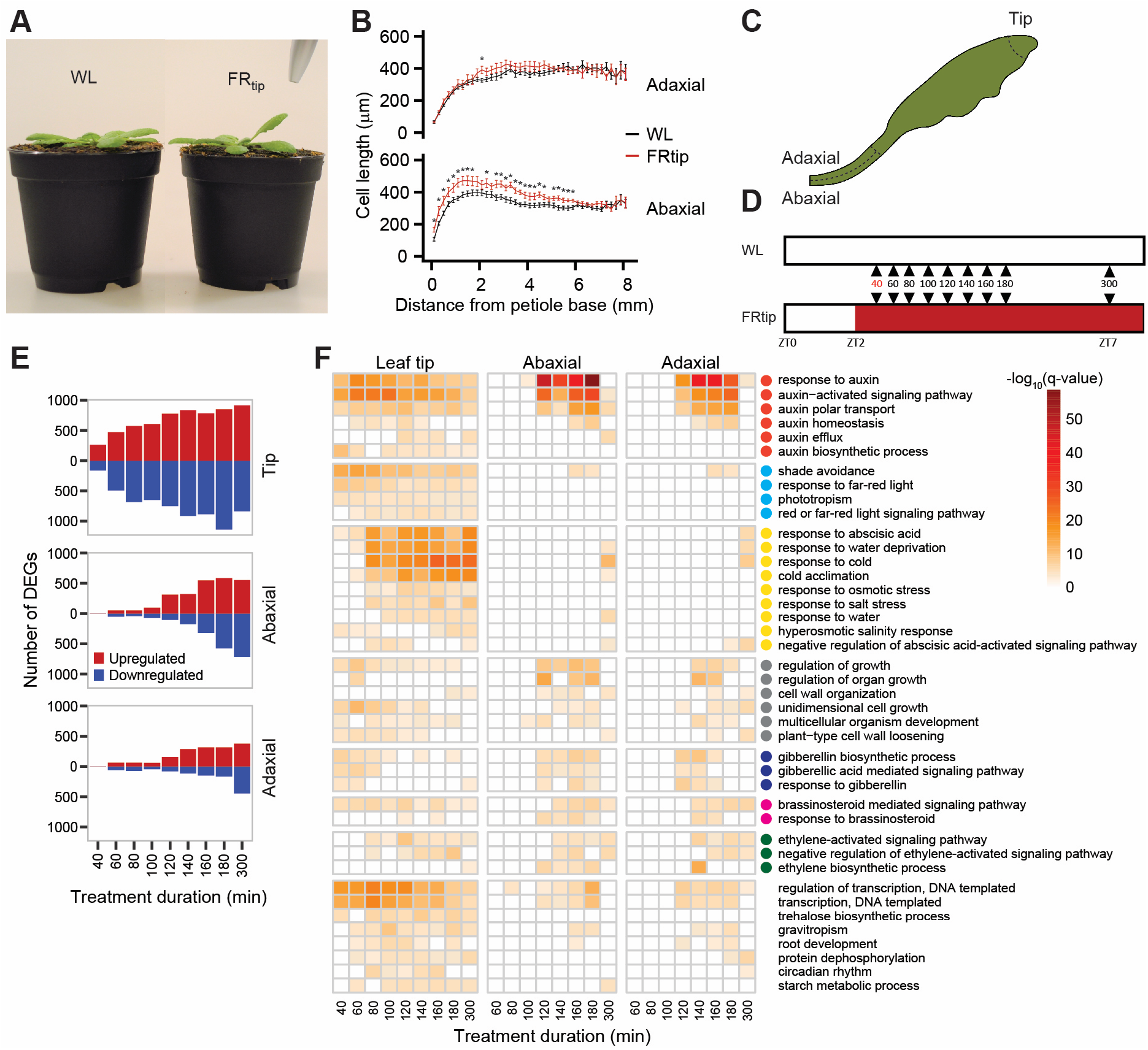
Neighbour detection in the leaf tip induces petiole hyponasty and transcriptional reprogramming in the petiole. (A) Adult Col-0 phenotype after 24h in the indicated light treatments. (B) Epidermal cell length measured along the abaxial and adaxial petiole after 24h in the indicated light treatments (n = 12 - WL, 15 - FRtip, *: p < 0.05, two-sided t-test, data represent mean ± SEM). (C & D) Schematic representations of harvested material (C, dotted lines identify the harvested sections in leaf tip and petiole base) and harvest timepoints (D) for RNA-sequencing. At the 40 min. timepoint, only leaf tip material was analysed. (E) Number of differentially expressed genes (DEGs) in FRtip compared to WL, calculated per timepoint and per tissue. DEGs were called when p < 0.01 and log_2_FC > 0.3 (upregulated; red) or log_2_FC < −0.3 (downregulated; blue). (F) Heatmap showing −log_10_(q-value) of gene ontology (GO) terms identified per timepoint and per tissue based on upregulated DEGs defined in (E). Coloured circles represent the following defined major biological processes; red – auxin distribution and signalling; cyan – light signalling; yellow – abscisic acid signalling; grey – cell and organ growth; blue – gibberellin biosynthesis and signalling; magenta – brassinosteroid signalling; green – ethylene biosynthesis and signalling. See also Figures S1 and S2, Video S1.

### Neighbour detection at the leaf tip induces local and remote, tissue-specific transcriptome changes

Reads were annotated to the TAIR10 genome and DESeq2-normalised read counts ^18^ were used to perform principal coordinate analysis (PCoA). We found clear PCoA separation between samples for timepoint and tissue type (Figure S1C). PCoA per tissue and differentially expressed gene (DEG) analysis per timepoint per tissue showed strong and consistent treatment effects in the leaf tip while in the petiole the treatment effect only became apparent at later timepoints (Figures 1E and S1D), consistent with our initial gene expression analyses (Figure S1B).

### Neighbour detection at the leaf tip induces tissue specific hormone response and biosynthesis

Gene ontology (GO) analysis for biological processes on upregulated DEGs per tissue per timepoint revealed early enrichment of auxin and light quality-related GO terms in the leaf tip followed by later enrichment of abscisic acid (ABA)-related GO terms (Figure 1F). As expected, light quality-related GO terms were largely absent from the petiole. In the petiole, we did, however, find enrichment of auxin response terms from 100 to 180 minutes, that dampened towards 300 minutes. This temporal GO enrichment pattern was similar for growth, response to brassinosteroid (BR) and ethylene as well as gibberellin biosynthesis and response (Figure 1F). Similar to the leaf tip, there was late enrichment of ABA-related GO terms in the petiole after the auxin response GO terms had passed peak significance. The apparent overrepresentation of auxin signalling in all tissues was confirmed when we analysed expression of all genes that make up the GO category GO:0009733 *“response to auxin”* (Figure S2A).

The analysis of these individual genes revealed shared, but also time and tissue-specific expression of many auxin-responsive genes. For example, regarding *SMALL AUXIN UPREGULATED* (*SAUR*) transcripts, *SAUR19-24* were induced in all tissues, while *SAUR25-29* and *SAUR62-68* were predominantly induced in the petiole (Figure S2A).

As we found GO enrichment for several hormone-related processes, we investigated expression of hormone biosynthesis genes (Figure S2B). Regarding the main auxin biosynthesis pathway, expression of *TRYPTOPHAN AMINOTRANSFERASE OF ARABIDOPSIS 1* (*TAA1*) and *YUCCA 6 (YUC6*) was repressed in the leaf tip while *YUC2*, *YUC5*, *YUC8* and *YUC9* expression was induced. In contrast, *YUC3* transcription was specifically induced in the petiole. Investigating gibberellin biosynthesis we found tissue-specific induction of *GA20 OXIDASE 1* (*GA20OX1*) and *GA20OX2* in the petiole and *GA20OX3* in the leaf tip. One step downstream of GA20OX proteins in the gibberellin biosynthesis pathway, *GA3 OXIDASE 1* (*GA3OX1*) was induced in both the leaf tip and the petiole. Regarding ABA biosynthesis, we found induction of *NCED3* in the leaf tip while *NCED5* was induced in the petiole. Besides auxin, gibberellin and ABA, we also observed transcriptional regulation of various genes involved in the biosynthesis of BR, ethylene and other hormones (Figure S2B).

To get a better insight in abaxial-adaxial transcript differences, we next identified genes that show differential response to FRtip between the two sides at 100 to 300 minutes of treatment (Figure 2). There were no genes with opposite regulation between the two sides but we did observe consistently stronger transcript regulation in the abaxial compared to the adaxial side of the petiole for both up- and downregulated DEGs (Figure 2A). The FRtip-upregulated genes in this subset showed enrichment for biological processes related to auxin and growth as well as to gibberellin, BR and ethylene (Figure 2B). As transcript regulation is strongest in the abaxial side of the petiole in this comparison this suggests that these processes are preferentially activated abaxially. Among the transcripts showing the highest significance in this analysis were many SAURs and other auxin-induced genes as well as the gibberellin biosynthesis genes *GA20OX1* and *GA20OX2*. Abaxial-adaxial transcriptional differences were also found in WL, and included many genes associated with photosynthesis.

**Figure 2.**
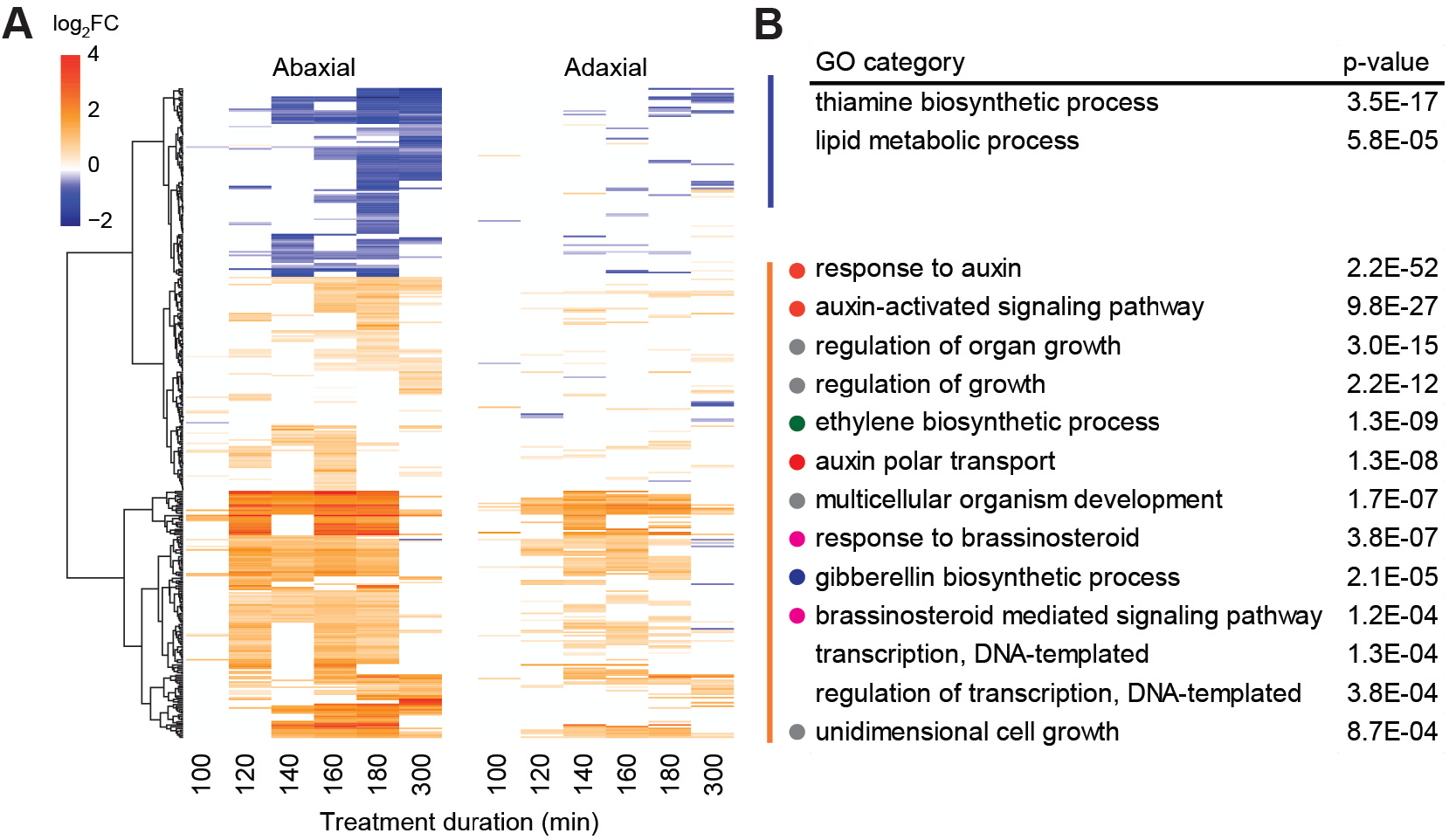
Neighbour detection at the leaf tip induces unique abaxial and adaxial transcriptomes. (A) Clustered heatmap showing log_2_FC in FRtip compared to WL of genes that show a different FRtip response between the two sides of the petiole at the indicated timepoints (ANOVA interaction tissue*treatment p < 0.001). (B) Separate GO analysis based on the clusters of upregulated (orange - red) and downregulated (blue) genes identified in (A). Coloured circles represent the following defined major biological processes; red – auxin distribution and signalling; grey – cell and organ growth; green – ethylene biosynthesis; magenta – brassinosteroid signalling; blue – gibberellin biosynthesis.

### Neighbour detection in the leaf tip leads to directed auxin transport towards the abaxial petiole

The enrichment for FRtip-induced auxin signalling in the transcriptome data prompted us to quantify free levels of the auxin indole-3-acetic acid (IAA) in the three leaf sections. We found increased IAA concentrations in the leaf tip and the abaxial petiole, but not in the adaxial petiole (Figure 3A) upon exposure of the leaf tip to FR. To study whether such differential auxin concentrations are required for petiole hyponasty, we exogenously applied IAA to the abaxial or adaxial petiole (Figure 3B). We found that abaxial IAA application results in strong hyponasty regardless of R/FR, while adaxial IAA application inhibited the hyponastic response to FRtip. These observations indicate that an auxin gradient, either installed endogenously or through directional external application, is necessary for leaf movement.

**Figure 3.**
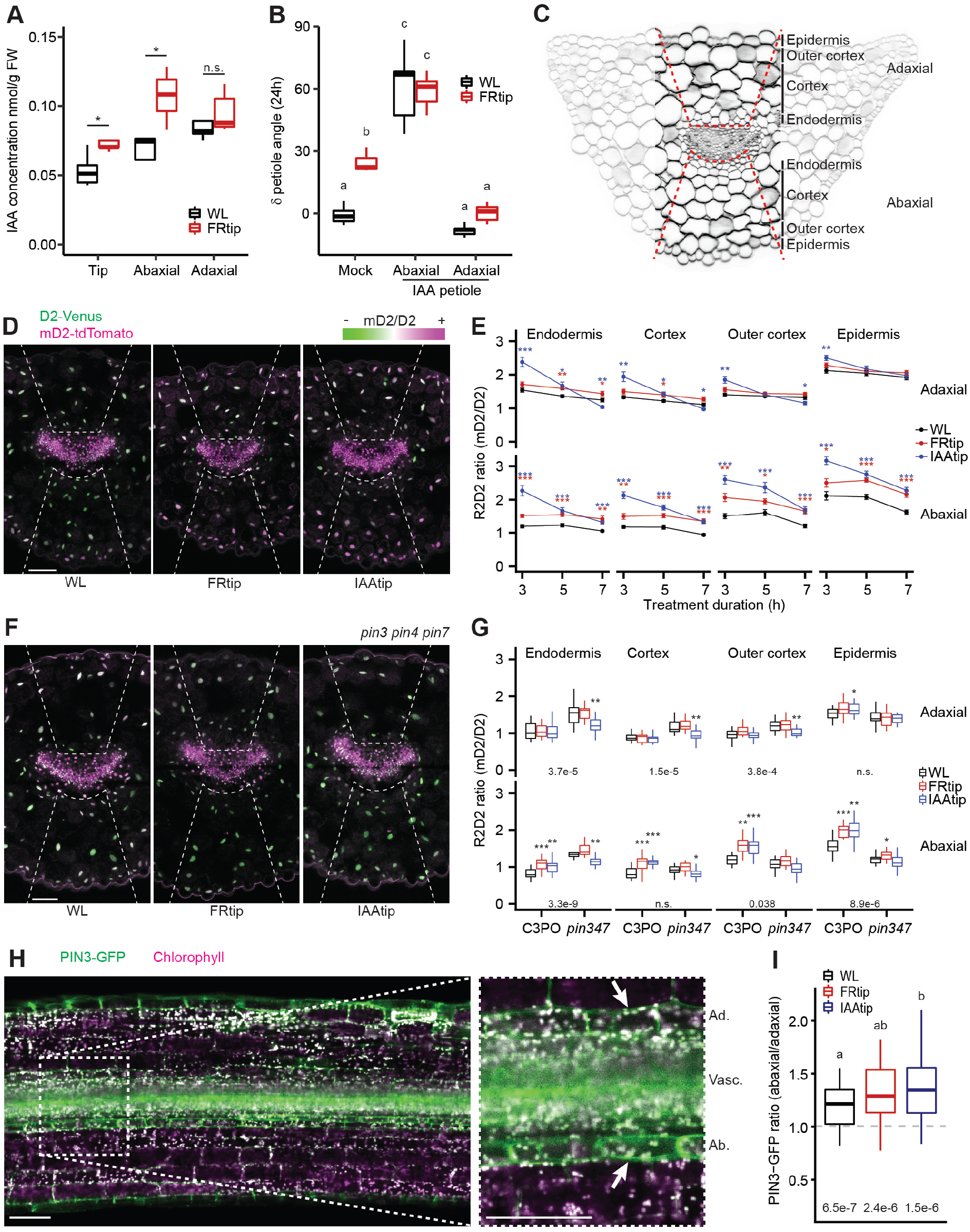
Leaf tip-derived auxin is directed towards the abaxial petiole via PIN transporters. (A) Free IAA concentration (nmol/g FW) in the leaf tip and abaxial/adaxial split petiole after 5h light treatment. (n = 5 biological replicates from 20 plants each, *: p < 0.05, two-sided t-test). (B) Petiole angle change after 24h light treatment combined with 30 μM IAA or mock application to the petiole. (n = 7, different letters indicate significant differences, Tukey HSD p < 0.05). (C) Petiole base cross-section indicating cell layers and region of the petiole that was used to quantify fluorescence in D – G and other figures. (D & E) Representative images after 5 h (D) and quantification at indicated timepoints (E) of the R2D2 ratio in the petiole base of C3PO. Plants were treated with mock, FRtip or IAAtip. (n = > 11, coloured asterisks represent significant treatment effect compared to WL, *: p < 0.05, **: p < 0.01, ***: p < 0.001, two-sided t-test, data represent mean ± SEM). (F & G) Representative images of *pin3 pin4 pin7* C3PO (F) and quantification of the R2D2 ratio in C3PO and *pin3 pin4 pin7* C3PO (*pin347*) (G) in the petiole base. Plants were treated for 7h with mock, FRtip or IAAtip. (n = > 15, asterisks indicate significant treatment effect compared to WL, *: p < 0.05, **: p < 0.01, ***: p < 0.001, two-sided t-test. Inset values represent p-value for genotype difference in WL calculated per cell layer, two-sided t-test). (H) Representative overview image and closeup around the vasculature of *pPIN3::PIN3-GFP* in a longitudinal petiole cross-section. Ad.: Adaxial endodermis, Vasc.: vasculature, Ab.: Abaxial endodermis. Arrows indicate the endodermal cells in which PIN3-GFP intensity in the membranes was quantified for I. (I) Ratio of PIN3-GFP intensity in the abaxial/adaxial endodermis after 2.5-4h in WL, FRtip or IAAtip. (n = 46 - WL, 28 - FRtip, 30 - IAAtip, different letters indicate significant differences, Tukey HSD p < 0.05). Inset values represent p-value for difference from ratio 1, one-sample t-tests. Scale bars in microscopy images represent 100 μm, dashed lines in D and F indicate the abaxial and adaxial regions where nuclear fluorescence was quantified. See also Figures S3 and S4.

To achieve further spatiotemporal resolution of auxin distribution, we visualised auxin distribution using the R2D2 part of the newly constructed C3PO fluorescent auxin reporter. C3PO conveniently combines the previously described R2D2 reporter for auxin concentration and the DR5v2 reporter that reports auxin response ^19^ into a single construct (*DR5v2::n3mTurquoise2-pRPS5A::mD2:ntdTomato-pRPS5A::D2:n3Venus*) (Figure S3). We developed a method to image transverse cross-sections of fixated and cleared petiole material in which we could measure fluorescence in individual cells and cell layers (Figures 3C and S3H). We found that the auxin concentration, as reported by the mD2/D2 (R2D2) intensity ratio, increased in all cell layers on the abaxial side within 3 hours of FRtip and remained higher than WL throughout the 7 hour interval that we measured, while there was little increase on the adaxial side (Figures 3D and 3E). When substituting FRtip with local IAA application on the leaf tip (IAAtip) we found increased R2D2 ratios after three hours in both sides of the petiole. At later timepoints of IAAtip treatment, the adaxial increase was lost and even changed into decreased R2D2 ratios in the adaxial endodermis and cortex, whereas the abaxial tissues continued to have an elevated R2D2 ratio, indicating elevated auxin levels.

### Auxin accumulation in the abaxial petiole via PINs

The petiole hyponasty response to FRtip requires intact auxin transport and is, therefore, reduced in the *pin3* single mutant and absent in the *pin3 pin4 pin7* triple mutant^10,11^. Similarly, *pin3* and *pin3 pin4 pin7* mutants respectively showed reduced and absent hyponasty in response to auxin application to the leaf tip (Figure S4A). When we analysed auxin distribution using the R2D2 ratio from C3PO crossed to the *pin3 pin4 pin7* mutant background we found that these mutations inhibited FRtip and IAAtip-induced abaxial R2D2 ratio increases (Figures 3F and 3G). The R2D2 ratio in WL was also different from wild type in *pin3 pin4 pin7* with a relatively increased R2D2 ratio in the inner cell layers and a reduced R2D2 ratio in the abaxial outer cortex and epidermis in *pin3 pin4 pin7* compared to wild type. We observed similar differences from wild type when regarding the auxin response, visualised by DR5v2::mTurquoise2 (Figures S4B and S4C), implying that perturbed PIN function prevents auxin transport towards the outermost cell layers in the petiole. In contrast with the induction of the R2D2 ratio by FRtip and IAAtip in C3PO, we did not find clear induction of DR5v2::mTurquoise2 intensity (Figures S4B and S4C). The lack of DR5v2 inducibility by IAAtip and FRtip likely indicates a poor sensitivity of this reporter in the petiole since our transcriptome analysis shows pronounced induction of auxin response upon FRtip (Figure 1), and only very large changes, such as following from the *pin3 pin4 pin7* triple mutant, affect the DR5V2 signal in petiole tissue.

Given the prominent effect of *pin* mutations on hyponasty (Figure S4A) and reported abaxial auxin accumulation in response to IAAtip and FRtip (Figure 3G), and the established regulation of PIN3 localization by supplemental FR in seedlings ^20^ we studied PIN3 localisation and abundance in petioles using *pPIN3::PIN3-GFP*. We found that in the petiole endodermis, PIN3-GFP is significantly enriched on the abaxial side compared to the adaxial side and that this asymmetry is reinforced in IAAtip treatment (Figures 3H and 3I). Taken together, this implies that PIN-dependent auxin transport directs tip-derived auxin to the abaxial petiole to stimulate abaxial cell elongation and petiole hyponasty upon neighbour detection in the leaf tip.

### Auxin activates members of the BAP/D module in the abaxial petiole for petiole hyponasty

Upon arrival in target tissue, auxin can stimulate growth by activating target gene expression via AUXIN RESPONSE FACTOR (ARF) transcription factors ^21^. Mutant phenotyping revealed that higher order mutant combinations of *ARF6, ARF7* (*NON-PHOTOTROPIC HYPOCOTYL 4, NPH4*) and *ARF8*, which were previously described to collectively regulate hypocotyl elongation responses ^22^, reduce the hyponastic response to tip-derived auxin (Figure 4A). ARF6 is one of the members of the BAP/D module, in which the transcription factors BZR1, ARF6 and PIF4 stimulate cell growth by reinforcing each other’s activity while all being repressed by DELLAs ^23^. PIF4, PIF5 and PIF7 together regulate FR-induced hyponasty ^11^ and mutation of *PIF4* and *PIF5* also reduced the petiole hyponasty response to IAAtip (Figure 4B). Loss of PIF7, in wild type or *pif4 pif5* background, however had little to no effect on the responsiveness to IAAtip. We observed a similar pattern when we applied IAA directly to the abaxial petiole (Figure 4C), confirming that the auxin response, and not auxin transport, is reduced in *pif4 pif5*, whereas *pif7* has a wild-type auxin response. Combined with our previous observation that FR-induced expression of *YUCCA* in the leaf tip is PIF7-dependent ^11^, we conclude that PIF7 is required for YUCCA-mediated auxin biosynthesis in the leaf tip, while PIF4 and PIF5 promote the auxin response in the petiole, probably as components of the BAP/D module.

**Figure 4.**
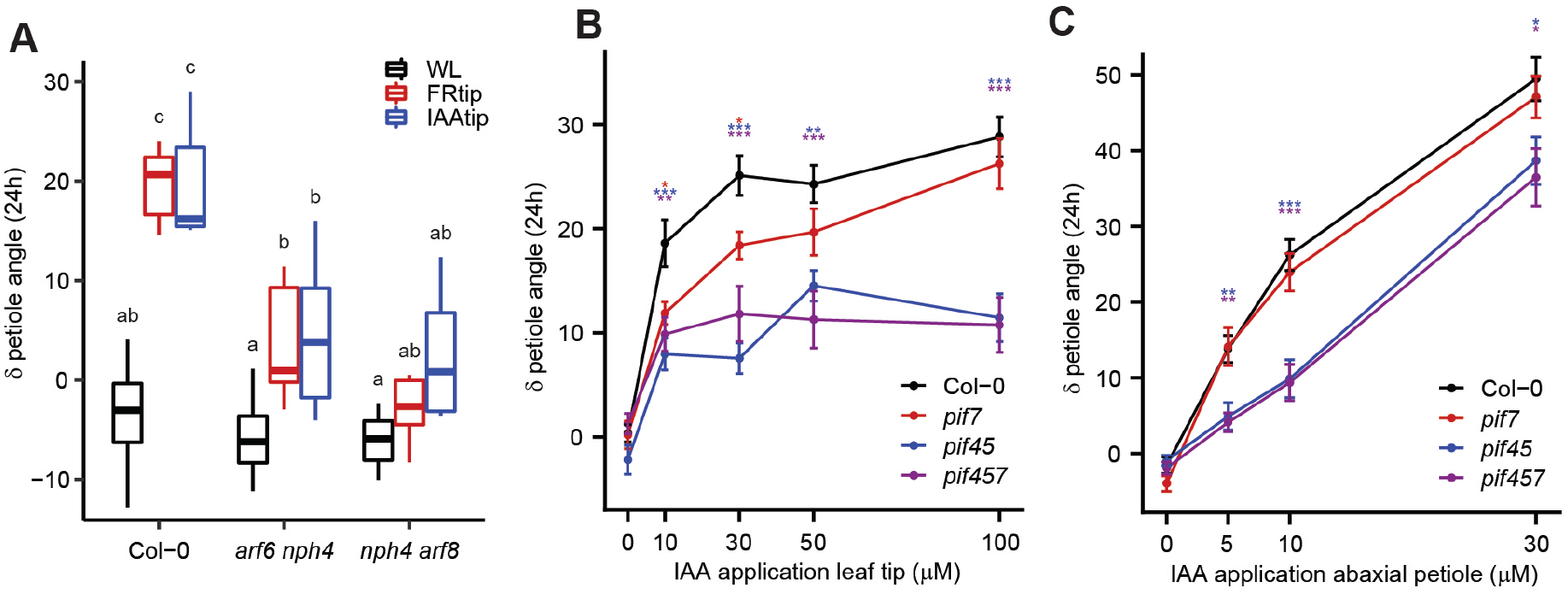
Leaf tip-derived auxin stimulates petiole hyponasty through activation of PIFs and ARFs. (A) Petiole angle change after 24h WL, FRtip or IAAtip treatment in Col-0, *arf6 nph4* and *nph4 arf8*. (n = 7, different letters indicate significant differences, Tukey HSD p < 0.05). (B & C) Petiole angle change after 24h in Col-0, *pif7, pif4 pif5* (*pif45*) and *pif4 pif5 pif7* (*pif457*)treated with different concentrations of IAA or mock to the leaf tip (B) and abaxial petiole (C). (n = 14, coloured asterisks represent significant genotype effect compared to Col-0, *: p < 0.05, **: p < 0.01, ***: p < 0.001, two-sided t-test, data represent mean ± SEM).

### Gibberellin as a downstream target of auxin signalling

The growth-repressing members of the BAP/D module, the DELLA proteins, are degraded through gibberellin signalling ^24^. In addition to the auxin enrichment profiles, our transcriptome analysis shows a strong enrichment for gibberellin biosynthesis and signalling, specifically in the abaxial petiole (Figures 1F and 2), where expression of *GA20OX1* and *GA20OX2* was induced (Figure S2B). This seems to be a response to tip-derived auxin as similar asymmetric induction of *GA20OX2* was found in response to IAAtip (Figure 5A).

**Figure 5.**
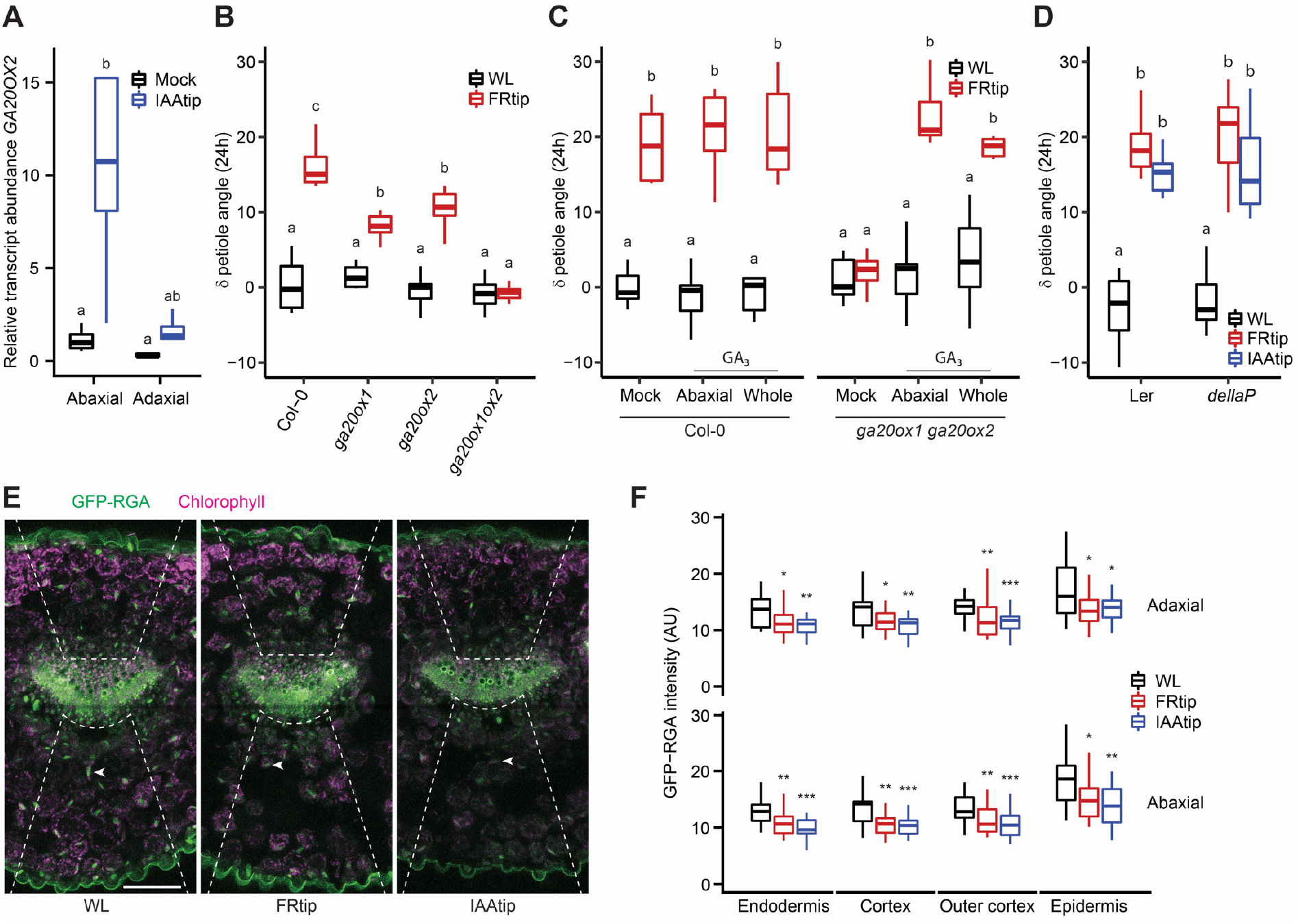
Gibberellin signalling facilitates the petiole hyponasty response to leaf tip-derived auxin. (A) Relative *GA20OX2* transcript abundance in the abaxial and adaxial petiole after 2h mock and IAAtip treatments. Relative transcript abundance compared to the abaxial petiole in mock treatment. (n = 4 biological replicates from 8 plants each, different letters indicate significant differences, Tukey HSD p < 0.05). (B) Petiole angle change after 24h light treatment in Col-0, *ga20ox1*, *ga20ox2* and *ga20ox1 ga20ox2* (*ga20ox1ox2*). (n = 9, different letters indicate significant differences, Tukey HSD p < 0.05). (C) Petiole angle change after 24h light treatment combined with 50 μM GA3 or mock application to the abaxial or whole petiole in Col-0 and *ga20ox1 ga20ox2*. (n = 7, different letters indicate significant differences, Tukey HSD p < 0.05). (D) Petiole angle change after 24h WL, FRtip or IAAtip treatment in *Ler* and *dellaP*. (n = 7, different letters indicate significant differences, Tukey HSD p < 0.05). (E & F) Representative images (E) and quantification (F) of GFP-RGA fluorescence in the petiole base. Plants were treated for 7h with mock, FRtip or IAAtip. (> 20, asterisks represent significant treatment effect compared to WL, *: p < 0.05, **: p < 0.01, ***: p < 0.001, two-sided t-test). Scale bar in E represents 100 μm, dashed lines indicate the abaxial and adaxial regions where nuclear GFP signal was quantified, arrowheads point out an individual nucleus in the abaxial cortex in each image. See also Figure S5 and Table S1.

Mutant analysis revealed that single *ga20ox1* and *ga20ox2* mutants showed reduced hyponastic responses to FRtip, and the *ga20ox1 ga20ox2* double mutant lacked all petiole hyponasty (Figure 5B).

When we applied GA to the petiole the hyponastic response to FRtip was restored in *ga20ox1 ga20ox2* (Figure 5C). Consistent with the mutant data, paclobutrazol (PAC) pre-treatment, which blocks gibberellin biosynthesis, also inhibited the hyponastic response to FRtip and this could also be rescued by exogenous GA application to the petiole (Figure S5A). In addition, we observed petiole hyponasty when we applied GA to the leaf tip in both wild type and *ga20ox1 ga20ox2* without additional FR (Figure S5B). Next, we tested the global (pentuple) DELLA knockout mutant *dellaP*.Although leaf angles were constitutively high in *dellaP*, FRtip and IAAtip still induced further petiole hyponasty, resulting in nearly vertical leaves (Figures 5D and S5C). When we studied DELLA abundance using the DELLA reporter *pRGA::GFP-RGA* that has been previously shown be GA-sensitive under low R:FR treatments, we observed clear RGA degradation in both sides of the petiole upon FRtip and IAAtip (Figures 5E and 5F). These data together indicate that leaf tip-derived auxin induces the expression of *GA20OX* gibberellin synthesis genes in the petiole, presumably leading to increased gibberellin levels. Indeed, leaf-tip derived auxin results in DELLA protein degradation in the petiole and the hyponastic response is gibberellin-dependent.

## Discussion

In this work, we show that plants use directional auxin transport from the leaf tip towards the abaxial petiole to initiate petiole hyponasty upon neighbour detection in the leaf tip. Using transcriptome analysis we reveal that phytochrome signalling of far-red light in the leaf tip induces a rapid auxin response in the abaxial petiole, that also stimulates expression of *GA20OX* gibberellin biosynthesis genes (Figures 1, 2 and S3).

The directed auxin transport towards the abaxial petiole requires functional PIN auxin efflux proteins (Figures 3F, 3G and S4), including PIN3. We show that in the petiole endodermis PIN3 is more abundant on the abaxial than the adaxial side and that this PIN3 asymmetry is enhanced in response to auxin application at the leaf tip (Figures 3H and 3I). The PIN3 asymmetry likely directs tip-derived auxin flow from the vasculature towards the abaxial petiole, thereby stimulating asymmetric cell growth and hyponasty (Figure 6). PIN4 and PIN7 localisation dynamics may also contribute to the directional auxin flow, potentially in other cell layers, as occurs in roots ^25^, but this was not investigated here. Endodermal PIN3 redistribution also occurs during FR light-induced hypocotyl elongation and during phototropism ^20,26^. Moreover, the petiole hyponastic response to elevated temperatures also involves PIN3 accumulation in the abaxial endodermis ^27^. However, these previously published examples involve direct light or temperature treatment exposure of the tissues where PIN3 redistributes. Our observation that remote IAAtip triggers similar endodermal PIN3 redistribution in the distal petiole, which is not exposed to treatment, implies that auxin itself reinforces the endodermal PIN3 asymmetry such that auxin is predominantly directed towards the abaxial side of the petiole. In support of this hypothesis, Keuskamp *et al*., 2010 showed that the FR-induced changes in PIN3 abundance and localisation in elongating hypocotyls relied on signalling of auxin itself. Possibly, the basic levels of auxin biosynthesis under control conditions suffice to create a basal level of PIN3 asymmetry in the petiole that is enlarged by additional FR-induced auxin biosynthesis. Other putative factors that could contribute to the abaxial-adaxial PIN3 asymmetry include signalling via leaf-polarity factors ^27–29^, asymmetric leaf and vasculature structure (Figure 3C), gravity ^30^ and even a light signalling gradient within the tissue ^31^.

**Figure 6.**
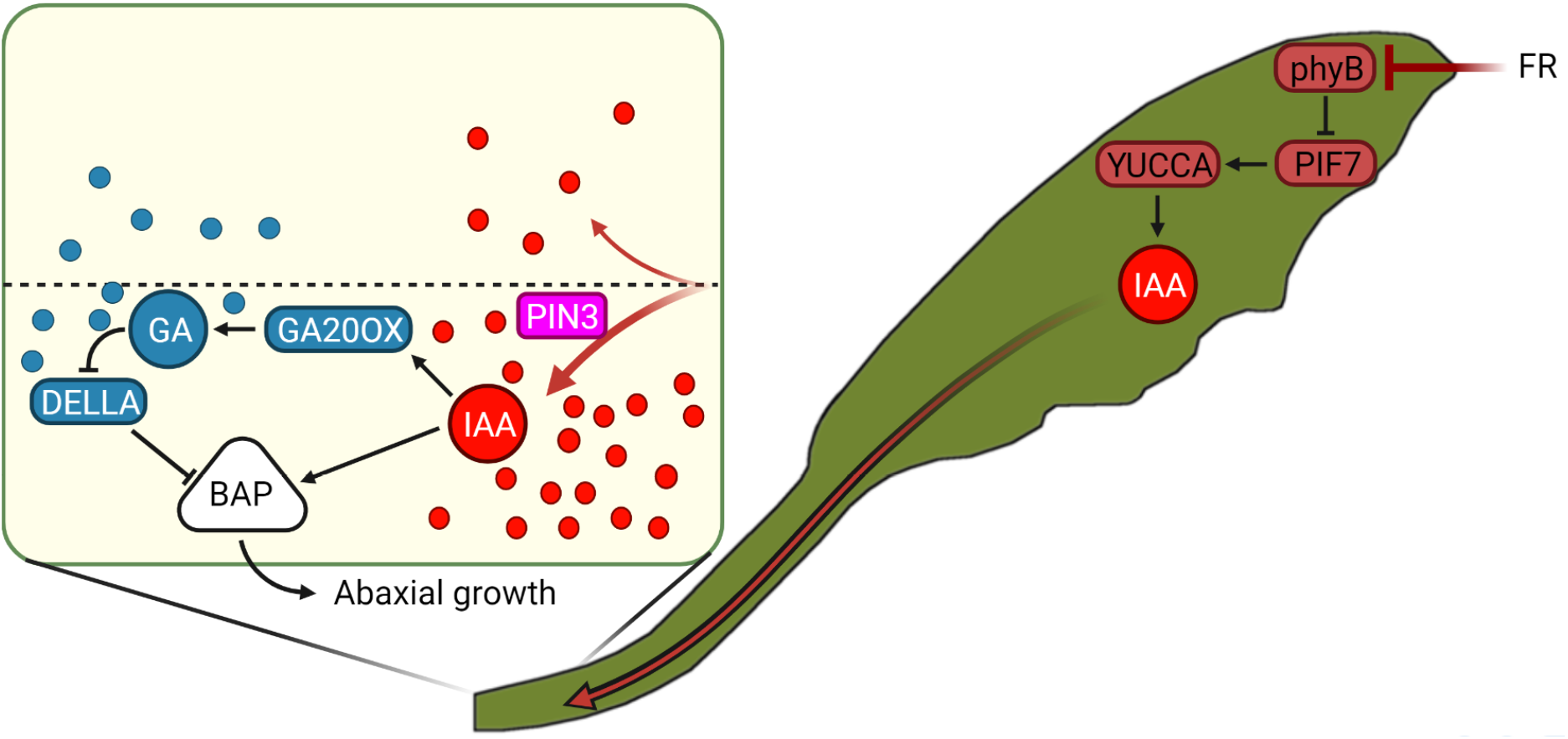
Proposed mechanism of how long-distance phytochrome signalling from tip to base orchestrates petiole hyponasty. FR light reflected from neighbours is first detected at the outermost leaf tip. This induces local inactivation of phyB, followed by auxin synthesis via PIF7 and YUCCAs, of which gene expression is induced within 40 minutes of FRtip. Auxin is transported from the leaf tip to the petiole and directed towards the abaxial petiole by PINs. In the abaxial petiole, leaf tip-derived auxin likely stimulates gibberellin synthesis via *GA20OX* expression, leading to the breakdown of DELLAs in the petiole. DELLA inactivation would then release repression of the auxin-activated growth-promoting BAP module. The asymmetric auxin distribution and signalling ensures that cell growth is limited to the abaxial petiole which results in adaptive petiole hyponasty. Round shapes represent auxin (IAA, red) and gibberellin (GA, blue).

Upregulation of *GA20OX* expression during shade and auxin-induced growth was previously shown ^32,33^. However, it was not known that these genes are also responsive to remote FR or auxin signalling. We observed that tip-derived auxin stimulates gibberellin biosynthesis by inducing *GA20OX1* and *GA20OX2* expression in the growing abaxial petiole and that as a result, the DELLA protein RGA is degraded (Figures 5A, 5E, 5F and S2B). In high gibberellin conditions, DELLA degradation prevents their inhibition of various growth-promoting transcription factors, including PIFs ^34,35^. In contrast to the specifically abaxial auxin accumulation and *GA20OX* expression, RGA degradation occurred non-specifically on both sides of the petiole (Figures 5E and 5F), suggesting abaxial-adaxial gibberellin transport that would result in non-differential gibberellin signalling in the petiole in response to FRtip. When we applied GA to both sides of the petiole in the gibberellin-deficient *ga20ox1 ga20ox2* mutant, we found that the hyponastic response to FRtip was rescued in a similar manner compared to when GA was applied only to the abaxial side and that GA application to the petiole in WL did not affect petiole angles (Figure 5C). We therefore propose that gibberellin abundance and subsequent DELLA degradation in the petiole are required to allow for petiole cell growth, while abaxial auxin accumulation provides the directional cue that ensures differential petiole growth that results in adaptive petiole hyponasty (Figure 6).

In contrast with *GA20OX1* and *GA20OX2*, a third member of the family *GA20OX3* was strongly induced specifically in the leaf tip by FRtip treatment. Our proposed mechanism for auxin-induced gibberellin biosynthesis in the petiole in a *GA20OX1* and *GA20OX2*-dependent manner, does not exclude the possibility that gibberellin derived from the leaf tip would also be transported towards the petiole to enhance petiole hyponasty. Indeed, when we applied GA to the leaf tip in WL, this resulted in petiole hyponasty in both wild type and *ga20ox1 ga20ox2* (Figure S5B). Keeping in mind that GA treatment of the petiole does not stimulate petiole hyponasty in WL (Figure 5C), we hypothesise that GA treatment of the leaf tip may locally degrade DELLAs, leading to enhanced PIF activity and auxin biosynthesis in the leaf tip. As *ga20ox1 ga20ox2* requires GA supplementation to the petiole to show petiole hyponasty in FRtip (Figure 5C), this suggests that exogenous GA to the leaf tip would also be transported towards the petiole. Combined tip-to-base transport of GA and auxin would then allow for hyponasty in *ga20ox1 ga20ox2*.

Besides their role in auxin biosynthesis, we showed that PIF4 and PIF5 are also required for the downstream petiole growth response to tip-derived auxin (Figures 4B and 4C). In addition, we found functional requirement for ARF6, ARF7 and ARF8 for petiole hyponasty, and transcriptional activation of BR signalling in the petiole (Figures 1F, 2 and 4A). This indicates that activation of members of the BAP/D module is involved in auxin-mediated petiole hyponasty (Figure 6). It remains to be studied whether the specific members and interactions in the BAP/D module are the same in adult petioles as in hypocotyls ^23^.

Spatial separation of light signalling and shoot growth response has been studied in seedlings in the past ^16,17,36^. However, the study system presented here provides an opportunity to study the effects of FR enrichment on distal, auxin-mediated growth without local light treatment of the responding organ. This will help further unravel the complex interactions between photoreceptors, the BAP/D module and other growth repressors and activators that plants use to optimize their growth to the environment ^37^. We conclude that upon neighbour detection, plants use carefully controlled long-distance auxin transport from the leaf tip to the abaxial petiole base to adaptively raise their leaves in a process that requires gibberellin biosynthesis and activation of the BAP/D module.

## Acknowledgements

We would like to thank all members of the Plant Ecophysiology research group at Utrecht University that participated in material harvests for RNA sequencing. We thank Kerstin Gühl for developing methods for IAA quantification and Yorrit van der Kaa for help propagating seed material. We also thank the Utrecht Sequencing Facility and the UMCU Bioinformatics Expertise Core for RNA sequencing and read annotation.

J.J.K. was supported by the Netherlands Organisation for Scientific Research (GSU 831.15.003), C.-Y.L. was supported by the European Research Council (ERC; StG “CELLPATTERN”; contract 281573 to D.W.), W.K. was supported by the Netherlands Organisation for Scientific Research (Veni 863.15.010), C.K.P. was supported by the Netherlands Organisation for Scientific Research (open competition ALWOP.509 to Kaisa Kajala), R.P. and L.O. were supported by the Netherlands Organisation for Scientific Research (Vici 865.17.002 to R.P.).

## Author contributions

Conceptualization, J.J.K. and R.P.; Methodology, J.J.K., L.O., S.E.A.M., C.-Y.L. and W.K.; Software, J.J.K. and B.L.S.; Formal Analysis, J.J.K. and B.L.S.; Investigation, J.J.K., L.O., C.K.P., S.E.A.M., E.R., E.D.C.E., H.W., C.-Y.L. and W.K.; Resources, C.-Y.L. and D.W.; Writing – Original Draft, J.J.K. and R.P.; Writing – Review & Editing, J.J.K. and R.P. with input from all authors; Visualization, J.J.K. and B.L.S.; Supervision, R.P.; Project Administration, R.P.; Funding Acquisition, J.J.K. and R.P.

## Materials and Methods

### Plant material and growth conditions

Genotypes used in this study: *ga20ox1-3* ^38^, *ga20ox2-1* ^38^, *ga20ox1-3 ga20ox2-1* ^38^, *arf6-2 nph4-1* ^22^, *nph4-1 arf8-3* ^22^, *pif4-101 pif5-1* ^39^, *pif7-1* ^40^, *pif4-101 pif5-1 pif7-1* ^41^, *pin3-3* ^42^, *pin3-3 pin4 pin7 ^43^, pin3-3 pPIN3::PIN3-GFP* ^44^, C3PO and *pin3 pin4 pin7* C3PO were all in Col-0 background; *dellaP* ^34^ and *pRGA::GFP-RGA* ^45^ were in L*er* background.

Seeds were sown on Primasta soil or agarose plates for germination and cold stratified for three days before transfer to short day white light (WL) conditions light/dark 9 h/15 h, 20 °C, 70 % humidity, 130-150 μmol m^-2^ s^-1^ PAR. Around eight days after germination, individual seedlings were transplanted to 70 mL round pots containing Primasta soil.

For microscopic screening of C3PO fluorescence in the root, seeds were surface sterilized, sown on half-strength Murashige and Skoog medium with 0.8% Daichin agar (Duchefa) (1/2 MS plate) and vernalized at 4 °C for 2 d. Afterwards, the seedlings were grown in climate room conditions at 22 °C in 16 h/8 h light/dark cycles.

### Construction of the C3PO auxin reporter

The C3PO construct (pGIIM/DR5v2::n3mTurquoise2-pRPS5A::mD2:ntdTomato-pRPS5A::D2:n3Venus) was generated via inserting DR5v2::n3mTurquoise2 into R2D2 ^19^. n3mTurquoise2 was generated by sequentially cloning the following three constructs, that were generated via PCR from plasmid template “pmTurquoise2-C1100”, into pGIIK/LIC_SwaI-LIC_HpaIv2-tNOS: mTurquoise2 coding sequence (CDS) with a stop codon, mTurquoise2 CDS without stop codon and NLS: mTurquoise2 without stop codon. The n3mTurquoise2-tNOS cassette was then excised via BamHI-XbaI double-digestion and inserted via conventional cloning into pGIIK/DR5v2::ntdTomato-tNOS, after the ntdTomato-tNOS cassette had first been removed via BamHI-XbaI double-digestion, to generate pGIIK/DR5v2::n3mTurquoise2-tNOS. An AscI restriction site was inserted into XbaI-digested pGIIK/DR5v2::n3mTurquoise2-tNOS via conventional cloning before ligating DR5v2::n3mTurquoise2-tNOS, that was excised by Bsp120I-AscI double-digestion, with Bsp120I-AscI double-digested pGIIM/pRPS5A::mD2:ntdTomato-pRPS5A::D2:n3Venus to generate pGIIM/DR5v2::n3mTurquoise2-pRPS5A::mD2:ntdTomato-pRPS5A::D2:n3Venus that we named C3PO. C3PO was then introduced into *Arabidopsis* via floral dip and selected using methotrexate. *pin3 pin4 pin7* C3PO was generated by crossing C3PO to *pin3-3 pin4 pin7*. Primer sequences used for cloning are shown in Table S1.

### Light and pharmacological treatments

For FRtip light treatment, WL was supplemented with FR using EPITEX L730-06AU FR LEDs. These FR LEDs had peak emission at 730 nm and locally reduced R/FR from ~2.0 in WL to below 0.1 in FRtip.

For pharmacological treatments at the leaf tip, 5 μL solution was pipetted onto the leaf tip. Except for the IAA concentration series in Figures 4B and S4A, 30 μM IAA was provided for IAAtip treatments. Pharmacological solutions and mocks for leaf tip application contained DMSO for IAA (0.03-0.1 %) or EtOH for GA_3_ (0.05 %) as well as Tween-20 (0.1 %). For hormone application to the petiole, concentrated stocks were diluted in lanolin (95-97 % lanolin, 0.01-0.03 % DMSO for IAA, 0.05 % EtOH for GA_3_). The lanolin containing solutions were carefully applied to the petiole using a tooth pick. When hormones were applied to one side of the petiole, a mock solution was applied to the other side. Paclobutrazol (PAC) treatment was done ten and five days before the experiment started. On both days, 20 mL 100 μM PAC or mock (0.3 % EtOH) was provided to the soil of each individual pot.

For all experiments, 28 day old plants were selected based on homogeneous development and the presence of a ~5 mm petiole on the 5^th^ youngest leaf which would be used in the experiment. All experiments were started at 10:00 (ZT2). For phenotyping experiments, petiole angle before treatment and after 24 hours was determined in ImageJ using side photos.

### Epidermal imprints and cell size measurements

Leaf material for epidermal imprints was harvested after 24 hours treatment. Dissected petioles were gently pressed into dental paste mixture (Coltene) to produce a leaf mold. After a few minutes of drying, a thin layer of transparent nail polish was applied onto the partially hardened dental paste before application of a second layer of dental paste on the adaxial side of the petiole. After solidification, the petiole sample was removed from the dental paste and a thin layer of transparent nail polish was brushed onto the imprint. The nail polish film was mounted on a microscopy slide and imaged at 40x magnification. Images were digitally stitched together and abaxial and adaxial cell lengths were measured along the petiole in ICY software (de Chaumont *et al*., 2012). Data was smoothened using a rolling average combining cell length data from up to 5 x-axis positions, depending on whether neighbouring datapoints were available.

### qRT-PCR and RNA-sequencing

For gene expression experiments, leaf tip and petiole material was harvested and snap frozen in liquid nitrogen and stored at −80 °C until further processing. The number of plants per replicate and number of replicates used in qRT-PCR experiments are indicated in the figure legends. RNA for qRT-PCR was isolated using the Qiagen RNeasy kit with on-column DNAse treatment. cDNA was synthesized using SuperScript III Reverse Transcriptase and random hexamer primers (Invitrogen). qRT-PCR was performed on the ViiA7 platform (Thermo Fisher) in 384-well plates using a 5 μL total volume containing SYBR Green (Bio-Rad). Transcript abundance was compared to housekeeping genes *PEX4* and *RHIP1* and made relative to the abundance in a designated control condition (indicated in figure legends). Primer sequences used for qRT-PCR are shown in Table S1. For RNA-sequencing, we harvested material from 13 leaves per sample, for a total of four biological replicates. Poly-A mRNA was isolated and used for the preparation of barcoded cDNA libraries according to the BrAD-seq protocol ^47^. Libraries were sequenced on an Illumina NextSeq 500 platform at 1*75bp read length yielding around 13 million reads per sample.

### RNA-sequencing data analysis

Reads were annotated to the TAIR10 genome and read counts were normalised using DESeq2 ^18^ (https://github.com/UMCUGenetics/RNASeq, https://github.com/UMCUGenetics/RNASeq#differential-expression-analysis). Genes that had an average of less than 1 annotated read per sample were removed. For the remaining 19663 genes, we calculated the mean read count as well as log_2_FC and p-value between treatments. Treatment-induced differentially expressed genes (DEGs) were identified per timepoint and per tissue when p < 0.01 and log_2_FC > 0.3 / < −0.3. For Figure S2, a log_2_FC cut-off of > 1 / < −1 was used. For Figure 2, we used an ANOVA approach to find genes with a significant (p < 0.001) two-way interaction Treatment*Tissue between the two petiole halves at timepoints 100 - 300 minutes. Principal coordinate analysis was performed on log_2_ transformed relative transcript abundance. Gene ontology (GO) enrichment analyses were performed using the hypergeometric test available in R. GO terms are only shown when highly significantly enriched in one sample (−log_10_(q-value) > 25) or consistently significantly enriched in five or more samples (−log_10_(q-value) > 5).

### IAA extraction and quantification by liquid chromatography-tandem mass spectrometry

For the extraction of IAA from *A. thaliana* petioles, ~40 mg of snap-frozen leaf material was used per sample. Tissue was ground to a fine powder at −80°C using 3-mm stainless steel beads at 50 Hz for 2*30 seconds in a TissueLyser LT (Qiagen, Germantown, USA). Ground samples were extracted with 1 mL of cold methanol containing [phenyl 13C6]-IAA (0.1 nmol/mL) as an internal standard as previously described ^48^. Samples were filtered through a 0.45 μm Minisart SRP4 filter (Sartorius, Goettingen, Germany) and measured on the same day. IAA was analyzed on a Waters Xevo TQs tandem quadruple mass spectrometer as previously described ^49,50^.

### Confocal microscopy

For confocal microscopy in transverse petiole cross-sections we harvested leaves into 24-well plates containing 4 % paraformaldehyde in PBS (pH 6.8) with Tween-20 (0.05 %). After vacuum incubation for one hour, leaves were washed three times for two minutes in PBS and stored for up to 24 h in PBS. Next, leaves were dried and placed in an Eppendorf tube containing warm agarose (3.5 %) and transferred to ice to solidify the agarose. Solid agarose plugs were sectioned to 250 μm slices using a Leica VT1000S vibratome. The first two slices from the petiole base (~0-500 μm) were discarded, and the next two (~500-1000 μm) were moved to 24-well plates containing ClearSee medium ^51^ and incubated for at least 7 days before microscopy. For Figure 3C, after the initial clearing, ClearSee was supplemented with Calcofluor white (0.01 %, 5 h), and rinsed afterwards with ClearSee. Longitudinal cross-sections for PIN3-GFP were made by hand, without prior fixation or clearing. Samples were directly placed with the cut edge onto a coverslip container (Lab-Tek) and immediately imaged. Sample drying was prevented by adding wet filter paper around the sample and covering the combination with a coverslip.

Confocal microscopy was largely performed on a Zeiss LSM880 system using a 25x glycerol objective. For C3PO we used the following laser and filters; mTurquoise2 – 458 nm laser, 467-500 nm filter, Venus – 514 nm laser, 525-550 nm filter, tdTomato – 561 nm laser, 571-629 nm filter. For PIN3-GFP we used; GFP – 488 nm laser, 501-548 nm filter, chlorophyll – 561 nm laser, 651-704 nm filter. For GFP-RGA we used; GFP – 488 nm laser, 510-525 nm filter, chlorophyll – 561 nm laser, 641-691 nm filter. Z-stacks were generated and combined into maximum intensity projections for nuclear fluorescence intensity measurements in ICY software ^46^. For PIN3-GFP, mean fluorescence intensity was measured in ICY on all sides of the visible endodermal cells in a single representative Z-layer. ICY was also used to select representative microscopy images and adjust brightness and contrast for improved clarity. Image adjustments were performed the same way between treatments.

For the development of C3PO, confocal microscopy on roots was performed on a Leica SP5II system using a 20x water-immersion objective with the following laser and filters; mTurquoise2 – 458 nm laser, 468-495 nm filter, Venus – 514 nm laser, 524-540 nm filter, tdTomato – 561 nm laser, 571-630 nm filter.

### Statistical analyses and data visualisation

Specific details on statistical analyses can be found in the figure legends. In multi-comparison analyses, we performed multi-factorial ANOVA with Tukey’s HSD post hoc correction. Elsewhere, we used two-sided t-test with p < 0.05 cut-off. Graphs and heatmaps were prepared in R and finetuned in Adobe Illustrator. The schematic model of signalling in Figure 6 was made in BioRender.

## Data availability

The raw RNA sequencing data generated in this study will be made publicly available in the National Center for Biotechnology Information (NCBI) Gene Expression Omnibus (GEO) upon publication. All data and biological materials are available from the corresponding author upon request.

## Legends for supplemental video and dataset files

**Video S1. Dynamics of FRtip-induced leaf movement, Related to Figures 1 and S1A**

Treatment duration in hours is indicated by the timer. Similar-sized leaves in the WL (left) and FRtip (right) treatments are indicated with an orange dot.

