## Supplemental information for "Local light signalling at the leaf tip drives remote differential petiole growth through auxin-gibberellin dynamics"

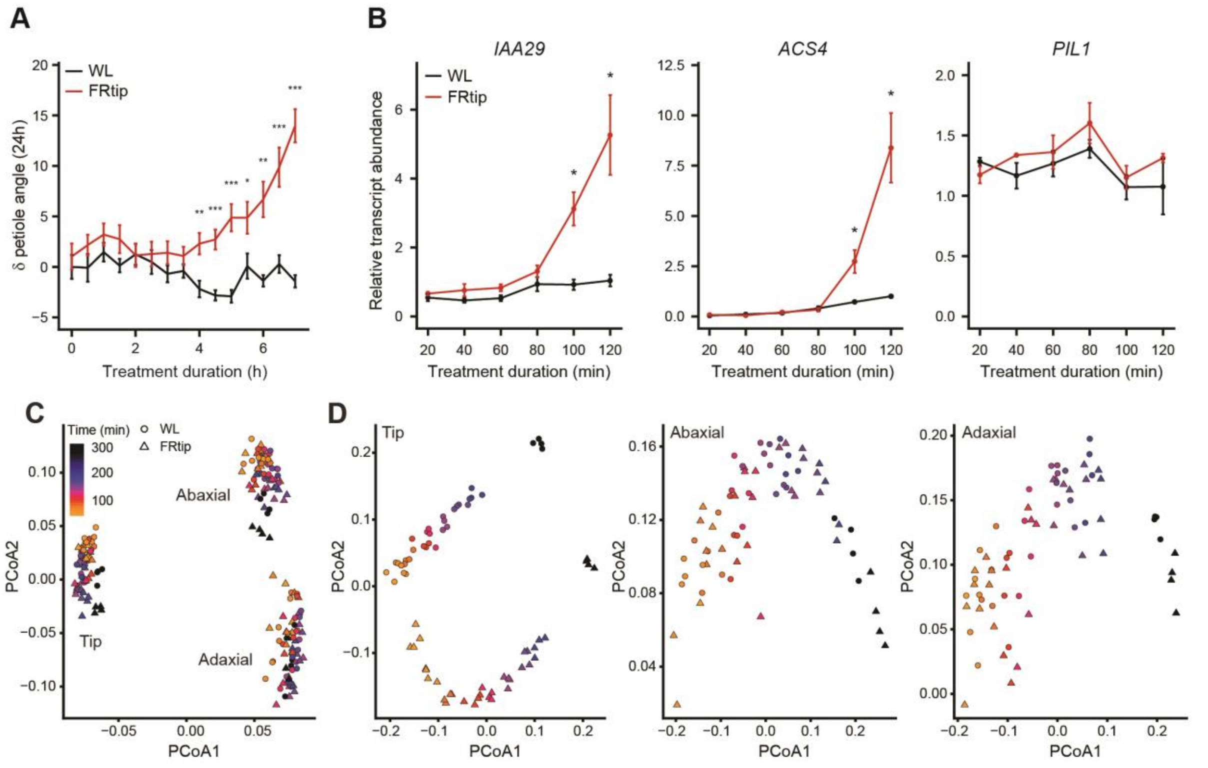


**Figure S1. Design criteria and principal coordinate analysis of RNA-sequencing experiment, Related to Figure 1**
(A) Petiole angle dynamics during the first 7 h of WL and FRtip. Petiole angle change is calculated relative to WL 0h treatment. (n = 10, *: $p < 0.05$, two-sided t-test per timepoint, data represent mean ± SEM). (B) Relative transcript abundance in the petiole base of the auxin response marker genes *IAA29* and *ACS4* and the shade marker gene *PIL1* during the first 2h of WL and FRtip. Transcript abundance is calculated relative to 120 min WL. (n = 2 (20 & 40 minutes), n = 4 (60 – 120 minutes), material harvested from 7 plants per sample, *: $p < 0.05$, two-sided t-test per timepoint, data represent mean ± SEM). (C & D) Principal coordinate analysis on all samples (C) and separately per tissue (D). See also Table S1.

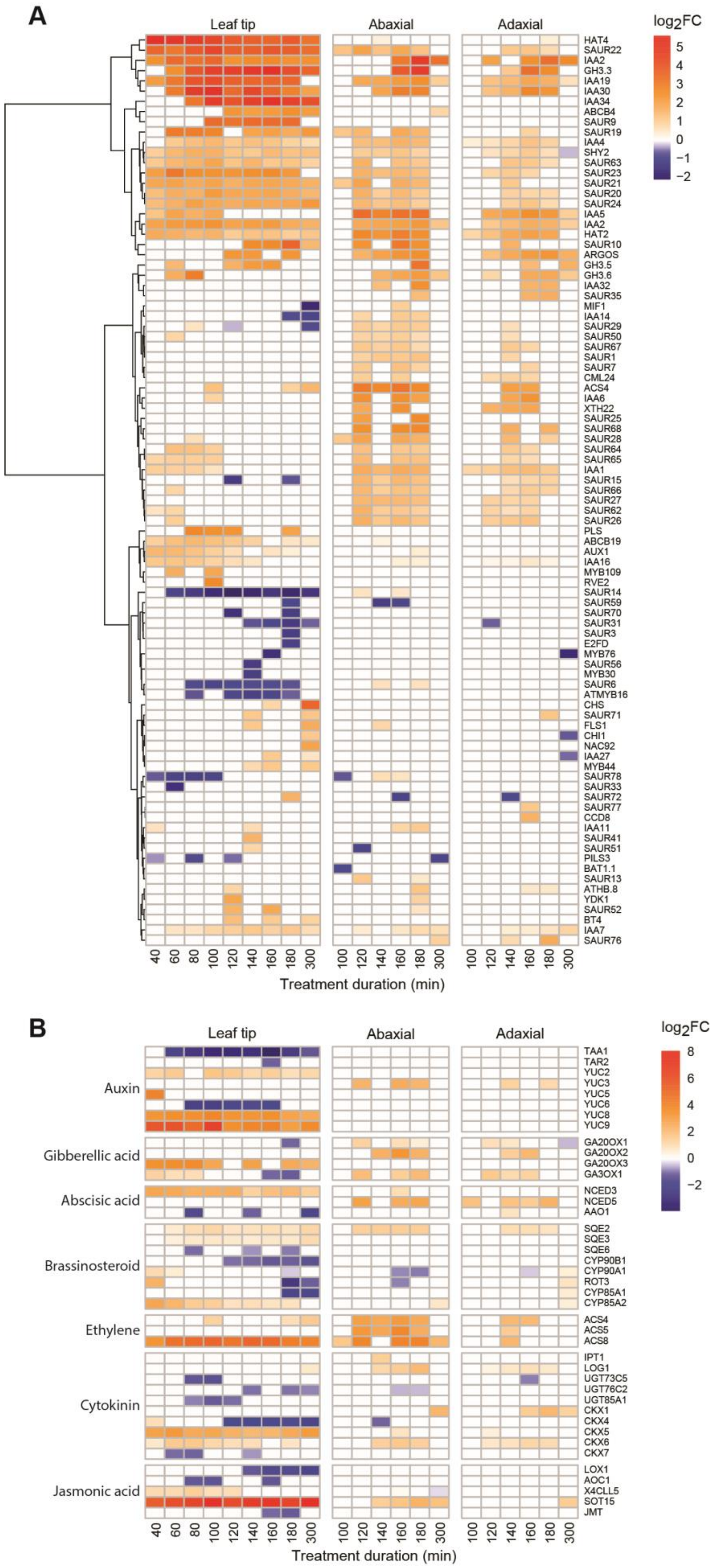
A
Leaf tip
Abaxial
Adaxial
log2FC
5
4
3
2
1
0
−1
−2
HAT4
SAUR22
IAA2
GH3.3
IAA19
IAA30
IAA34
ABCB4
SAUR9
SAUR19
IAA4
SHY2
SAUR63
SAUR23
SAUR21
SAUR20
SAUR24
IAA5
IAA2
HAT2
SAUR10
ARGOS
GH3.5
GH3.6
IAA32
SAUR35
MIF1
IAA14
SAUR29
SAUR50
SAUR67
SAUR1
SAUR7
CML24
ACS4
IAA6
XTH22
SAUR25
SAUR68
SAUR28
SAUR64
SAUR65
IAA1
SAUR15
SAUR66
SAUR27
SAUR62
SAUR26
PLS
ABCB19
AUX1
IAA16
MYB109
RVE2
SAUR14
SAUR59
SAUR70
SAUR31
SAUR3
E2FD
MYB76
SAUR56
MYB30
SAUR6
ATMYB16
CHS
SAUR71
FLS1
CHI1
NAC92
IAA27
MYB44
SAUR78
SAUR33
SAUR72
SAUR77
CCD8
IAA11
SAUR41
SAUR51
PILS3
BAT1.1
SAUR13
ATHB.8
YDK1
SAUR52
BT4
IAA7
SAUR76
40 60 80 100 120 140 160 180 300
100 120 140 160 180 300
100 120 140 160 180 300
Treatment duration (min)
B
Leaf tip
Abaxial
Adaxial
log2FC
8
6
4
2
0
−2
Auxin
Gibberellic acid
Abscisic acid
Brassinosteroid
Ethylene
Cytokinin
Jasmonic acid
TAA1
TAR2
YUC2
YUC3
YUC5
YUC6
YUC8
YUC9
GA20OX1
GA20OX2
GA20OX3
GA3OX1
NCED3
NCED5
AAO1
SQE2
SQE3
SQE6
CYP90B1
CYP90A1
ROT3
CYP85A1
CYP85A2
ACS4
ACS5
ACS8
IPT1
LOG1
UGT73C5
UGT76C2
UGT85A1
CKX1
CKX4
CKX5
CKX6
CKX7
LOX1
AOC1
X4CLL5
SOT15
JMT
40 60 80 100 120 140 160 180 300
100 120 140 160 180 300
100 120 140 160 180 300
Treatment duration (min)

**Figure S2. Neighbour detection at the leaf tip induces tissue-specific auxin response and hormone biosynthesis, Related to Figure 1**
(A) Clustered heatmap showing $log_2FC$ in FRtip compared to WL of genes in the GO category "GO:0009733 response to auxin" calculated per timepoint and per tissue. (B) Heatmap showing $log_2FC$ in FRtip compared to WL of genes involved in major hormone biosynthesis pathways calculated per timepoint and per tissue. In both heatmaps, only genes that have a significant treatment effect in at least one sample are shown ($p < 0.01$, $log_2FC > 1$ / $< -1$).

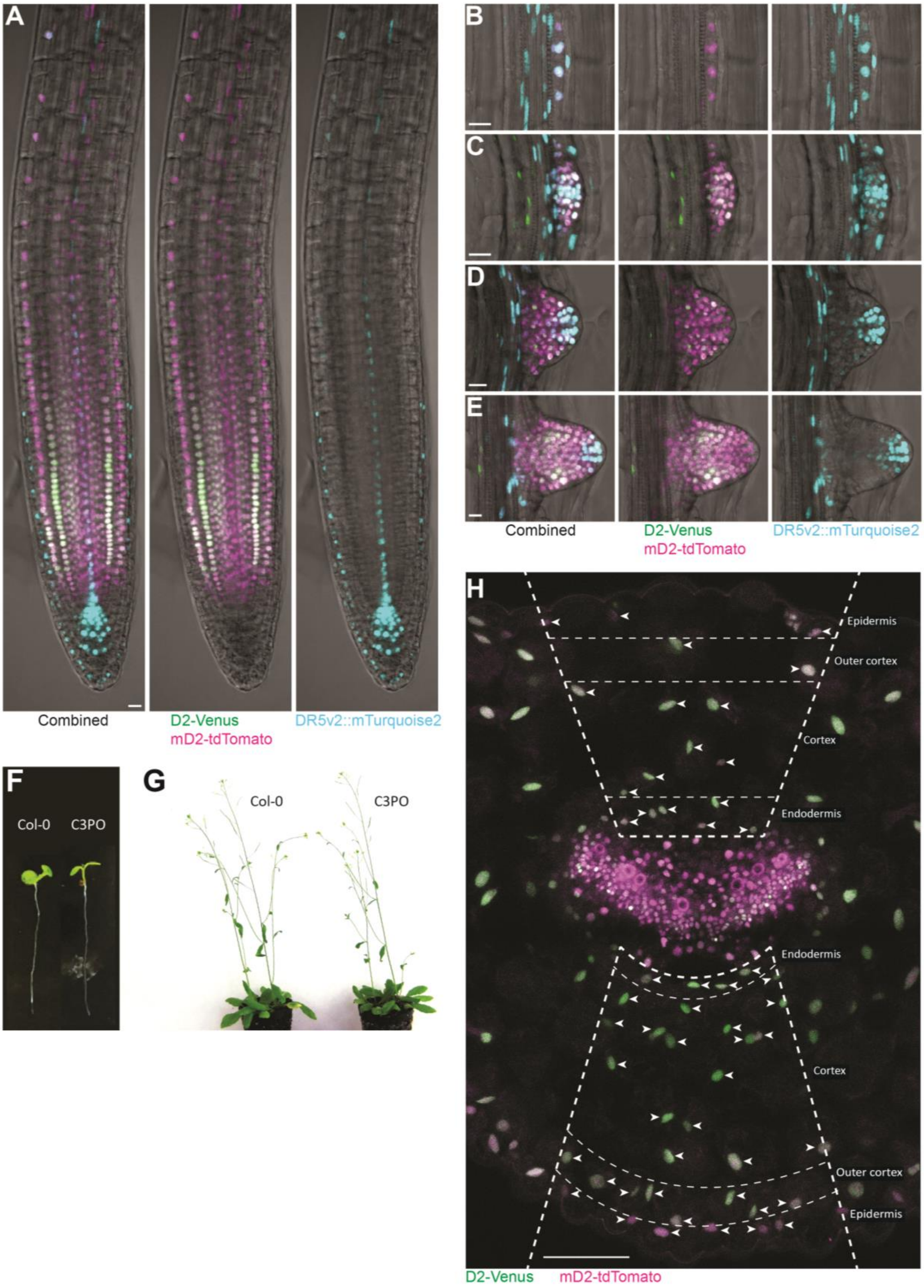


**Figure S3. Description of C3PO fluorescence patterns and intensity quantification, Related to Figure 3**
C3PO fluorescence patterns in the root tip (A), and lateral root primordium at stage I (B), stage IV (C), stage VI (D) and stage VIII (E) of 5 day old seedlings. (F & G) Plant phenotype of 6 day old seedling (F) and 35 day old flowering plant (G) of Col-0 and C3PO grown in 16 h/8 h light/dark conditions. (H) Close-up image of the transverse petiole cross-section of C3PO in WL (Figure 3D), showing D2-Venus and mD2-tdTomato fluorescence. Dashed lines indicate the region and cell types where C3PO fluorescence was imaged. Arrowheads indicate individual nuclei in which fluorescence intensity was quantified. Note that in the adaxial epidermis, stomatal fluorescence was not measured. Scale bars represent 10 µm in A – E and 100 µm in H. See also Table S1.

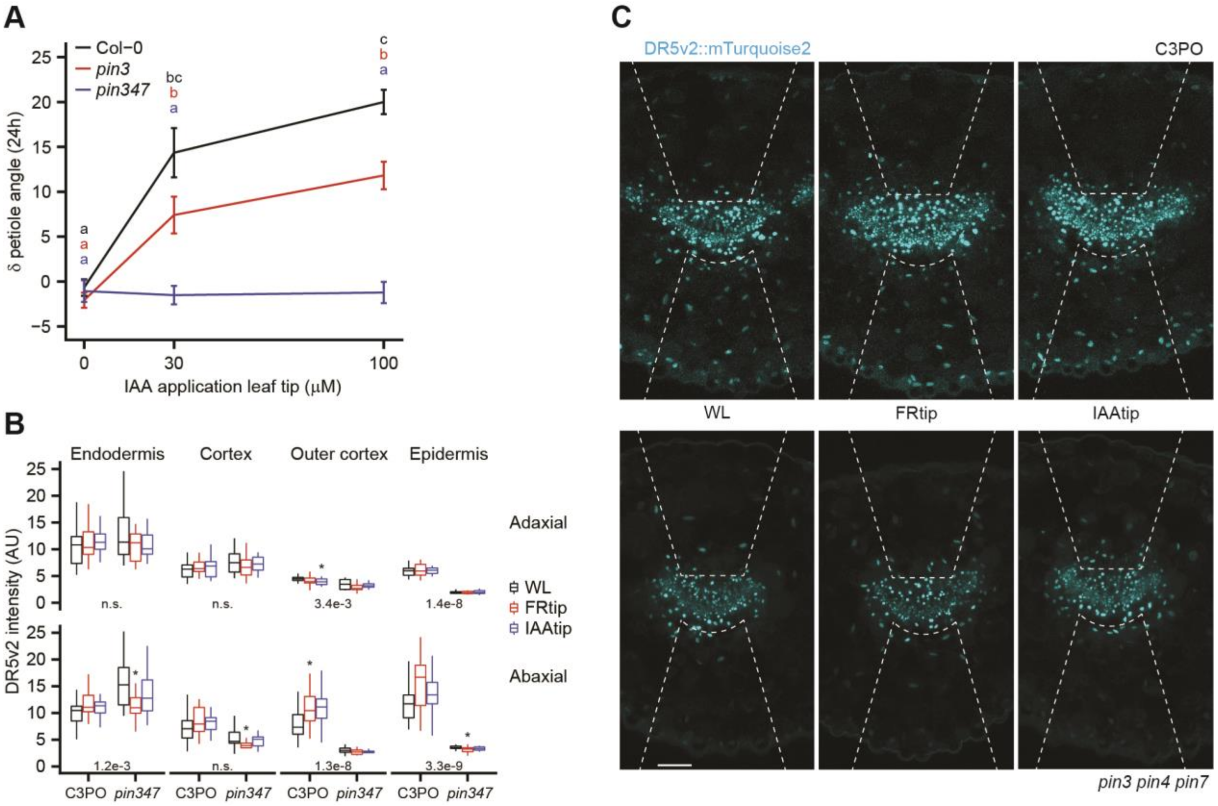


**Figure S4. PIN auxin transporters are required for auxin signalling in the petiole and the hyponastic response to leaf tip-derived auxin, Related to Figure 1**

(A) Petiole angle change after 24 h in Col-0, *pin3* and *pin3 pin4 pin7* (*pin347*) treated with different concentrations of IAA or mock to the leaf tip. (n = 10, different letters indicate significant differences with colours representing the corresponding genotype, Tukey HSD $p < 0.05$, data represent mean ± SEM). Representative images (B) and quantification (C) of DR5v2::mTurquoise2 fluorescence in the petiole base in C3PO and *pin3 pin4 pin7* C3PO. Plants were treated for 7 h with mock, FRtip or IAAtip. (n = > 15, asterisks indicate significant treatment effect compared to WL, *: $p < 0.05$, two-sided t-test. Inset values represent p-value for genotype difference in WL calculated per cell layer, two-sided t-test). Data from the same experiment as shown in Figures 3F and 3G. Scale bar represents 100 µm, dashed lines indicate the abaxial and adaxial regions where nuclear mTurquoise2 fluorescence was quantified.

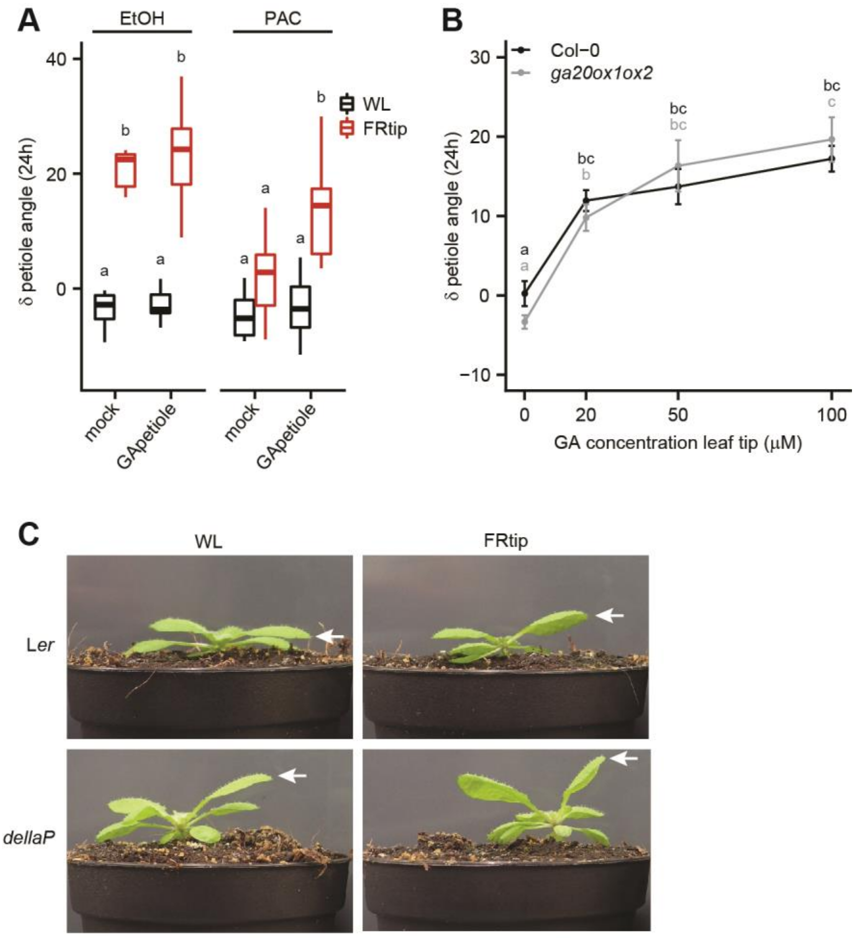


**Figure S5. Gibberellin signalling stimulates petiole hyponasty, Related to Figure 5**
(A) Petiole angle change after 24 h in mock (EtOH) or PAC pre-treated Col-0 plants treated with WL or FRtip and mock or 50 µM $GA_3$ to the abaxial petiole. (n = 7, different letters indicate significant differences, Tukey HSD $p < 0.05$). (B) Petiole angle change in Col-0 and *ga20ox1 ga20ox2* (*ga20ox1ox2*) after 24 h mock or $GA_3$ treatment to the leaf tip in WL conditions. (n = 7, different letters indicate significant differences with colours representing the corresponding genotype, Tukey HSD $p < 0.05$, data represent mean ± SEM). (C) Representative images of L*er* and *dellaP* after 24 h in WL and FRtip. The leaves of interest are indicated with the arrows.

**Table S1. Primers used in this study, Related to Figures 5A, S1 and S3**

| Primer set | Forward | Reverse | Purpose |
|---|---|---|---|
| ACS4 | GGAGCCACTTCCGCAAAC | GCTTGCTCGTAGGCTTCTTC | qPCR |
| IAA29 | AAGATGGATGGTGTGGCAAT | GTCACCCTCTTTCCCTTGGA | qPCR |
| PIL1 | AGACCACCTACGATGTTGCC | TAGCATTTGTGGTGGTGCAT | qPCR |
| GA20OX2 | AGTAGCTTCACCGGCAGATT | ACGCCTAAACTTAAGCCCAGA | qPCR |
| RHIP1 | ATTGGTGTCGCTGCTAGTCT | TAAAGCCGTCCTCTCAAGCA | qPCR |
| PEX4 | TGCAACCTCCTCAAGTTCGA | TGAGTCGCAGTTAAGAGGACT | qPCR |
| mTurqoise2 + STOP | TTTTGGATCCGGTGGTATGGT GAGCAAGGGCGAGGA | TTTTAGATCTTTACTTGTACA GCTCGTCCATGC | Cloning |
| mTurqoise2 non-STOP | TTTTGGATCCGGTGGTATGGT GAGCAAGGGCGAGGA | TTTTAGATCTCTTGTACAGCT CGTCCATGCC | Cloning |
| NLS-mTurqoise2 non-STOP | TTTTGGATCCCATGGCTCCAA AGAAGAAGAGAAAGGTCATG GTGAGCAAGGGCGAGGA | TTTTAGATCTCTTGTACAGCT CGTCCATGCC | Cloning |
| Additional AscI | CTAGATTAATTAAGACACAGG CGCGCCT | CTAGAGGCGCGCCTGTGTCT TAATTAAT | Cloning |
